# Interspecies differences in lamina cribrosa fiber composition and architecture might arise from differences in optic nerve head size and mechanical constraints

**DOI:** 10.64898/2026.11.18.767671

**Authors:** Adam E. Galloy, Xuehuan He, Ian A. Sigal

**Affiliations:** Department of Ophthalmology, University of Pittsburgh School of Medicine, Pittsburgh, Pennsylvania, USA; Department of Bioengineering, University of Pittsburgh, Pittsburgh, Pennsylvania, United States

**Keywords:** lamina cribrosa, intraocular pressure, eye movements, topology optimization, ocular biomechanics

## Abstract

The lamina cribrosa (LC) is a complex network of fibers that plays an important mechanical role in protecting optic nerve axons as they exit the eye. Both the architecture and composition of the LC network varies throughout animal species. However, the function of these differences between species is unknown. We hypothesize that this variety in LC architecture and composition is a result of species-specific adaptations to the mechanical constraints present in each species’ optic nerve head. To test this hypothesis, we developed a novel optimization framework to systematically derive optimal LC fiber architecture and composition under different mechanical constraints. We then compared predicted LC architectures to instant polarized light microscopy images of actual human, tree shrew, monkey, cow, pig, and rat LCs. Distinct features from the various LCs were replicated by varying the mechanical constraints, consistent with our hypothesis. The wheel-and-spoke architecture of the tree shrew LC was captured with angularly symmetric loading conditions. Non-radial fibers like those found in monkey and human LCs were captured by adding horizontal eye rotation tractions. The softer glial LC of the rat and stiffer collagenous LC of larger species were replicated by a two-material extension of the model. To our knowledge, these are the first results that systematically show how interspecies differences in LC architecture and composition can arise from mechanical constraints and size. This work can result in better interpretation of animal models and provide a point of comparison for changes in LC architecture under pathological conditions such as glaucoma.

## Introduction

The lamina cribrosa (LC) is a complex network of fibers at the optic nerve head—the opening in the sclera where retinal ganglion cells axons exit the eye on their way to the brain. The LC fibers form a porous network spanning the scleral canal near its opening [1–3]. These fibers provide structural integrity to the optic nerve head and protect retinal ganglion cell axons. When the LC’s protection is insufficient, axons can suffer from damage that ultimately leads to loss of vision [4–7]. Two well-accepted sources of load acting on the optic nerve head that the LC provides protection for are the intraocular pressure [4] and cerebrospinal fluid pressure [8]. Work over the last decade has shown that tractions produced by rotations of the eye can also significantly deform the LC and surrounding tissue in human eyes [9,10].

Between species there is a wide variety of LC architectures. Tree shrew LCs exhibit a wheel and spoke architecture with collagen fibers extending radially from the lamina to the central retinal vessel whereas monkey LCs and human LCs have more complex architectures of intercrossing and non-radially oriented fibers [11–15]. The LC also exhibits compositional variation between species. Many large mammal species—including humans, monkeys, tree shrews, and pigs—have an LC composed of collagen fibers (collagenous lamina) whereas in other species such as mice and rats the actin and glial fibrillary acidic protein form a similar fibrous network, forming the so-called *glial lamina* [11,16,17]. This wide array of architectures and compositions remains unexplained, betraying a lack of understanding on the relationship between LC structure and function. <u>We hypothesized that the interspecies variety in LC architecture and composition can be explained as unique adaptations to the mechanical constraints experienced by each species’ optic nerve head.</u> For example, the radially symmetric wheel-and-spoke architecture of the tree shrew LC suggests it has adapted to radially symmetric loading conditions (such as IOP) rather than directionally biased tractions (like tractions from eye rotations). Similarly, species with small eyes potentially do not require complex collagenous LCs because the stresses resulting from IOP are small enough to be handled with softer compositions.

To test our hypothesis, we developed a topology optimization model of the optic nerve head to determine how different loading conditions and geometric constraints impact optimal collagen architecture and composition. Topology optimization algorithms seek to find the distribution of material within a domain that minimizes material volume while maintaining structural integrity against a set of loads. Although topology optimization is typically used for the design of artificial constructs, it has proven useful when applied to biological tissues as well. For example, topology optimization has been used as a model for bone growth and remodeling after surgical intervention [18] and during aging [19]. The application of topology optimization to etiological and comparative biology studies is also well-established. Zhao et al. used topology optimization to relate differences in plant stalk cross-sections between species to the differences in mechanical conditions these species adapted to [20]. Sakashita et al. used topology optimization with similar goals in their study of fish vertebrae [21]. One thing shared by these studies on biological tissue was that the tissue of interest behaved like typical bulk engineering materials such as metal that can have arbitrary thickness and sustain compressive, shear, and tensile loads allowing for the adaptation of traditional topology optimization algorithms. In contrast, the LC is composed of slender fibers that primarily function in tension and have the tendency to buckle under compression, bending, or shear stresses [22,23]. Optimizing fibrous structures like the LC consequently requires a reformulation of the optimization problem and solution algorithm.

We therefore decided to base this project on the methods introduced by Rowe and Chern for the optimization of truss structures, which have negligible shear and bending stresses like slender fibers [24], but we introduced a new tension-only constraint to make the method completely suitable for optimizing fibrous biological tissue structures. We compared models of the optic nerve head optimized under various loading and geometry constraints to images of several animal species in order to systematically probe how these constraints could influence interspecies variations in LC architecture. To understand interspecies variations in LC composition, we extended the topology optimization model to include two different fiber materials. We used the two material model to test the hypothesis that a softer glial lamina may be appropriate to protect small eyes even when otherwise identical conditions in large eyes would require a stiffer collagenous lamina.

## Methods

We developed a novel topology optimization problem formulation and algorithm that enabled us to derive optimal fiber architectures from different mechanical conditions. To verify the implementation of the algorithm’s different features, we solved three two-dimensional example models. More complex three-dimensional optic nerve head models were then implemented within the topology optimization framework to conduct three systematic studies on LC architecture. First, the role of algorithm parameters was assessed on baseline models featuring a circular scleral canal and only the loads arising from pressures acting on the optic nerve head. Second, the role of scleral canal geometry was assessed by optimizing models with the same loading conditions as before, but with elliptical canals of varying aspect ratio. Third, the role of eye rotation tractions was assessed by adding horizontal shear tractions to the posterior surface of the LC models and solving with varying magnitudes of the shear tractions. To be able to use the algorithm to predict optimal LC composition in addition to the architecture, the optimization framework was extended to include two fiber materials: a weak material representing astrocytic actin and a strong material representing collagen. The role of animal size on optimal LC composition was assessed with models of small eyes and large eyes solved with otherwise identical loading and geometry conditions. Finally, instant polarized light microscopy (IPOL) images of human, tree shrew, monkey, cow, pig, and rat LCs were obtained to compare actual LC architectures to the architectures the models predicted under different conditions.

### Topology optimization overview

The optimization process minimizes the volume of fiber material required to bear a given load. Fibers are the main load bearing component of the LC. The fibers are long and slender with very small resistance to bending, compression, and shearing (assumed to be negligible inside of fibers for this work). Consequently, the optimization process needed to address the challenge of identifying an LC architecture that can resist bending, compression, and shear loads using constitutive elements that act primarily in uniaxial tension.

Briefly, the approach to solving this problem was to encode the spatial distribution, orientation, and loading state of fibers at a location with the stress tensor at that location. This represented the placement of fibers directly into the loaded configuration at a level of stretch consistent with their stress. The constitutive behavior of the LC fibers was captured via constraints applied directly to the stress tensor. The slenderness of the LC fibers was captured by requiring that the stress tensor was nearly everywhere uniaxial (a sparse set of elements where fibers can merge/branch was acceptable) and was nowhere in a state of compression. To prevent the fibers from being placed in overly distended and ineffectual states an upper bound on the tensile stress was added. The volume of fibers was then minimized by driving the stress tensor to zero on as much of the domain as possible while still satisfying static equilibrium and constitutive constraints. A more precise mathematical description of the problem and constraints is provided in the Topology optimization problem formulation section.

Since we assumed in this work that the fiber architecture was optimized for a particular loading condition, the fiber material properties and mechanics only needed to be considered in that loaded configuration. This eliminated the need to make assumptions about the unloaded configuration of the fibers (which need not be identical for each fiber [25]) and avoided numerical issues that are often present in stiffness-based formulations of slender structures. Interestingly, this approach produced architectures insensitive to the precise stiffness of the fibers since a softer/stiffer fiber could hypothetically still be placed at the same stress, but with greater/lesser stretch. However, for extreme differences in fiber stiffness (such as between actin and collagen fibers) the maximum allowable distension would occur at significantly lower stress in softer materials which must be captured by a different upper bound on stress. This is described in more detail for the two-material models in the Two-material models of the optic nerve head section.

To numerically solve the optimization problem, we developed a custom MATLAB script adapting Rowe and Chern’s algorithm for optimizing truss structures [24]. The algorithm optimized a nonconvex cost function by means of gradient descent with static equilibrium constraints enforced by the method of augmented Lagrangians. Full details on the numerical solution are provided in the Numerical implementation section.

### Topology optimization problem formulation

Following Rowe and Chern’s formulation [24], the optimization variable is the stress tensor field ***σ*** throughout the optimization domain Ω subject to the static equilibrium constraint:

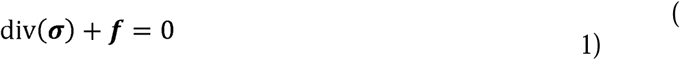

Where ***f*** is a vector field of body forces applied to the domain. The cost function *W* seeks to find ***σ*** with minimal support over the domain (i.e. ***σ*** = 0 on as much of the domain as possible) while encouraging all but one eigenvalue of ***σ*** to vanish in locations where the stress is nonzero:

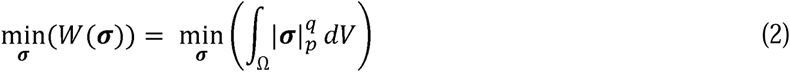

Where in terms of the eigenvalues of the stress tensor *σ*_1_, *σ*_2_, and *σ*_3_:

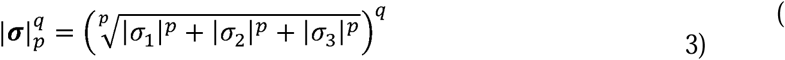

Exponential coefficients 0 < *p*, *q* < 1 result in a concave cost function that promotes sparsening of ***σ*** and its eigenvalues (see Rowe and Chern [24] for more details). The resulting sparse stress field ***σ*** encodes the optimized architecture. Locations where ***σ*** has nonzero components represent locations with material and locations where ***σ*** is (or is nearly) zero represent voids without material. The orientation of ***σ***’s nonzero eigenvector(s) indicate the orientation of the structural member(s) crossing that location.

The static equilibrium constraint on ***σ*** was enforced via the augmented Lagrangian method. The Lagrangian formulation of the constrained optimization problem is given by:

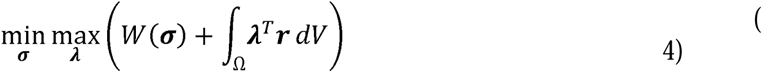

Where ***λ*** is the vector field of Lagrange multipliers and ***r*** is the residual of the static equilibrium constraint:

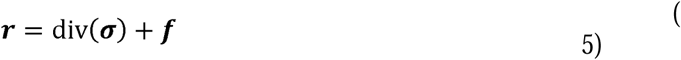

The augmented Lagrangian method approximates the true Lagrangian as follows:

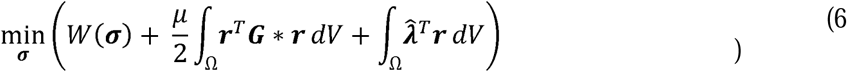

Where 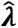 are approximations of ***λ***, *μ* is a penalty scaling constant, and ***G*** ∗ represents convolution of ***r*** with a regularization metric. The regularization metric ***G*** and approximation scheme for 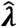 are provided in the *Numerical implementation* section.

Expanding on Rowe and Chern’s original problem formulation, two additional constraints were required for physically meaningful fiber architectures. First, biological fibers have negligible bending stiffness and buckle under slight compressive loads, so the eigenvalues of the stress tensor cannot be negative. Second, fibers have an upper limit on the stress that they can carry *σ_max_*. Without this limit the structures predicted can be unreasonably sparse resulting in highly concentrated loads that would compromise the LCs’ ability to protect the surrounding nerve axons. This gives the box constraints:

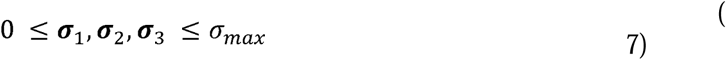

We found that the upper bound constraint could be strictly enforced without numerical difficulty. However, initial guesses for the stress field often contained many elements in compression. These compressed elements required large, unstable corrections to strictly enforce the lower bound. To alleviate these numerical issues, instead of strictly enforcing the lower bound we defined separate tensile and compressive cost functions:

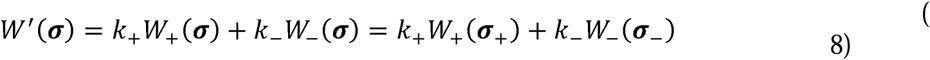

Where *k*_+_ and *k*_−_ are weights associated with each cost function component. The stress tensor field is decomposed into tensile and compressive components:

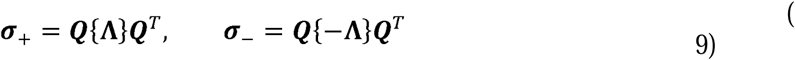

Where ***Q*Λ*Q****^T^* is the eigenvalue decomposition of ***σ*** and { } are Macaulay brackets. The tensile component of the cost function remains the same as the original cost function:

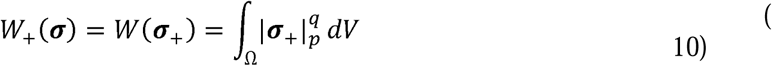

For our models we always used 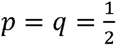 for the exponential coefficients. For the compressive component, an L1 penalty was chosen for the cost function:

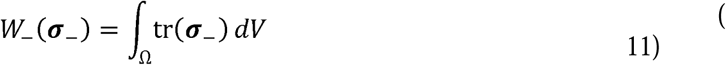

The L1 penalty is an exact penalty in the sense that, if a compression-free solution exists, there is some finite value of the penalty weight *k*_−_ that will result in the elimination of ***σ***_−_ [25]. By incrementally increasing the penalty weight, numerical issues can be avoided while allowing the solution to approach strong enforcement.

### Numerical implementation

The domain of the optimization problem was a rectangular prism with dimensions L_x_ x L_y_ x L_z_ discretized into a grid of N_x_ x N_y_ x N_z_ voxels. To allow for the use of Fourier analysis in solving the static equilibrium condition, the domain was assumed to be periodic. Obstacles (described later) were placed around the boundaries of every model to ensure this did not lead to unphysical structures.

The augmented Lagrangian problem is iteratively solved using a truncated gradient descent method. The constraint function, its residual, and its gradient are computed in Fourier space where the divergence operator and its adjoint can be represented as sparse matrices for fast and simple computation as presented in Eqs. (12(12)-(19). The cost function gradients are given in Eqs. (20-(23). From these gradients, the precise updating procedure for the optimization variables is presented in Eqs. (24-(28) and summarized in Algorithm 1. Implementation of the algorithm was done in MATLAB R2024b (The Mathworks Inc., Natick, MA).

#### Fourier representation of the static equilibrium constraint and its gradients

The divergence operator for a given combination of spatial frequencies {*k_x_*, *k_y_*, *k_z_*} for a symmetric tensor quantity in Voigt notation [*a*_11_, *a*_22_, *a*_33_, *a*_23_, *a*_13_, *a*_12_]^T^ is given by:

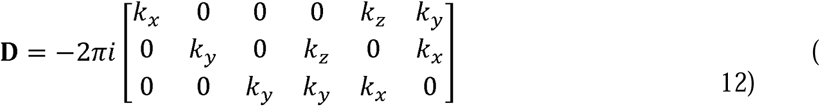

And its adjoint by:

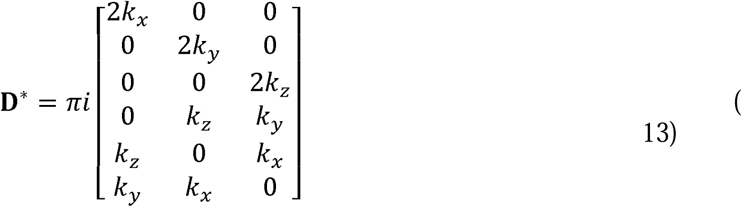

The residual function of the static equilibrium constraint is then given by:

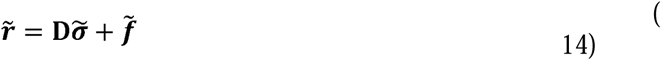

Where a tilde indicates the discrete Fourier representation of a quantity. The terms of the augmented Lagrangian problem Eq. (6) associated with the static equilibrium constraint become:

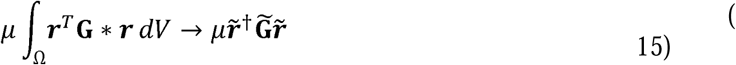

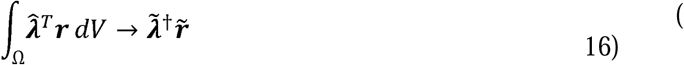

The operator **D** is unbounded with respect to frequency. To avoid ill-conditioning the following regularization metric ***G̃*** was used [24]:

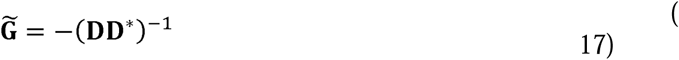

After substituting Eq. (14) and Eq. (17), the gradient of Eq. (15) with respect to stress is given in the Fourier domain by:

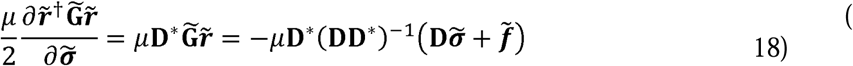

And the gradient of Eq. (16) is given by:

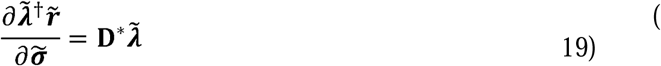

For all models we chose *μ* = 1.

#### Gradients of the cost function

Choosing 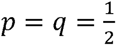 the gradient of the tensile cost function Eq. (10) is computed on each voxel after eigenvalue decomposition:

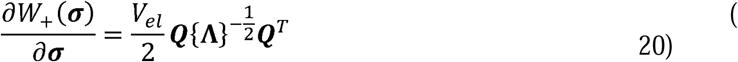

Where *V_el_* is the element volume. For zero eigenvalues where the gradient is undefined, we note that *W*_+_(***σ***) is already minimized with respect to that eigenvalue and take the gradient to be zero. The gradient of the compressive cost function Eq. (11) after eigenvalue decomposition is:

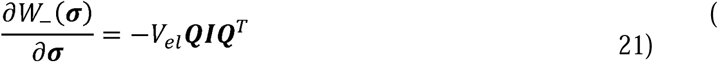

Where ***I*** is the identity tensor. The total gradient of Eq. (8) is then:

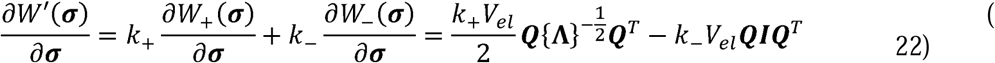

The magnitude of the weight parameters *k*_+_ and *k*_−_ play important roles in how quickly the optimization model converges and how accurately the constraints are enforced. To make identifying a value of *k*_+_ more physically intuitive, we relate it to another parameter called the critical stress *σ_cr_*:

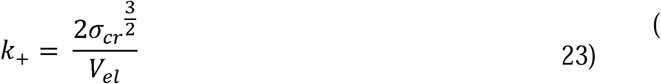

*σ_cr_* has the same units as stress. *σ_cr_* is the critical point where the scaled gradient of *W*_+_ with respect to a tensile stress eigenvalue equals that eigenvalue. Tensile eigenvalues near or below *σ_cr_* will have large gradients relative to their value and vanish quickly, whereas tensile eigenvalues above *σ_cr_* will have small gradients and are more likely to persist in the optimization process. Due to its more physically meaningful interpretation we report *σ_cr_* rather than *k*_+_ for each of our models. *k*_−_ was allowed to vary throughout the optimization process.

#### Iterative updating procedure

At every optimization iteration three quantities are updated: the optimization variable ***σ***, the approximate Lagrange multipliers ***λ̃***, and the compressive cost weight *k*_−_. All these quantities are set to zero at initialization. ***σ*** is updated in two steps, one in the Fourier domain, and another in its eigenbasis in the spatial domain. In the first step, the Fourier transform of ***σ*** at the current iteration, ***σ̃***_(*n*)_, and its static equilibrium residual ***r̃***_(*n*)_ are evaluated and a constraint step is taken according to the gradients of the constraint equations:

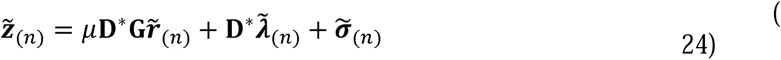

The inverse Fourier transform is then taken of ***z̃***_(*n*)_. In the second step, eigenvalue decomposition on ***z***_(*n*)_ is performed at each voxel. A truncated gradient step is then taken from the diagonal matrix of eigenvalues **Λ**_(*n*)_. The eigenvalues in tension are updated according to:

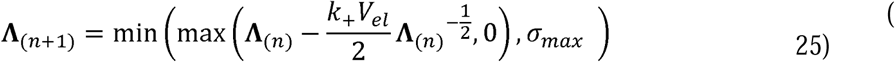

Where ***σ****_max_* is the maximum allowed tension. Substituting (23) one finds that tensile eigenvalues at or below *σ_cr_* are set to zero by this step. The eigenvalues in compression are updated according to:

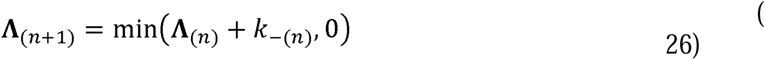

With *k*_−(*n*)_ determined by:

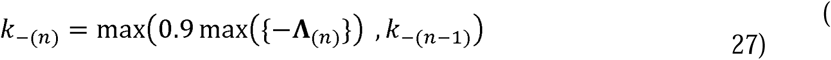

Strong enforcement is achieved when *k*_−(*n*)_ is greater than the largest compressive eigenvalue max({−**Λ**_(*n*)_}). Although the update rule in Eq. (27) appears to always choose a smaller value than this, we note that *k*_−(*n*)_ is only allowed to increase. As the compressive component is penalized and the algorithm prioritizes tensile members, max({−**Λ**_(*n*)_}) eventually starts decreasing and *k*_−(*n*)_ is large enough to satisfy the constraint. The updated eigenvalues are rotated back into the cartesian basis to get the updated optimization variable ***σ***_(*n*+1)_. The residual of the static equilibrium condition is computed in the Fourier domain for the updated stress field and the Lagrange multipliers are then updated according to:

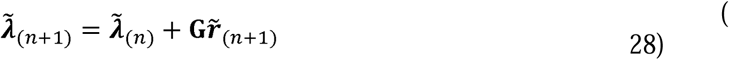

This process repeats a set number of iterations. The optimization loop is summarized in Algorithm 1.

#### Topology optimization inputs and outputs

Figure 1a summarizes the inputs and outputs accepted by the topology optimization model. Loads are prescribed elementwise and determine the static equilibrium constraint, Eq. (5). Pre-existing support structures for the optimized fiber architectures to build off are implemented by setting *k*_+_ and *k*_−_ to zero at elements belonging to the support structure. In other words, static equilibrium updates are performed on the stress tensor in these elements, but the stress tensor in these elements does not contribute to the cost function and is not changed by the eigenvalue updates (Eqs. (25) and (26)). Obstacles may also be assigned to elements in the domain, so that the algorithm does not place material there. This is implemented by scaling *k*_+_ and *k*_−_ by a factor of 10^6^ at these elements.

**Figure 1:**
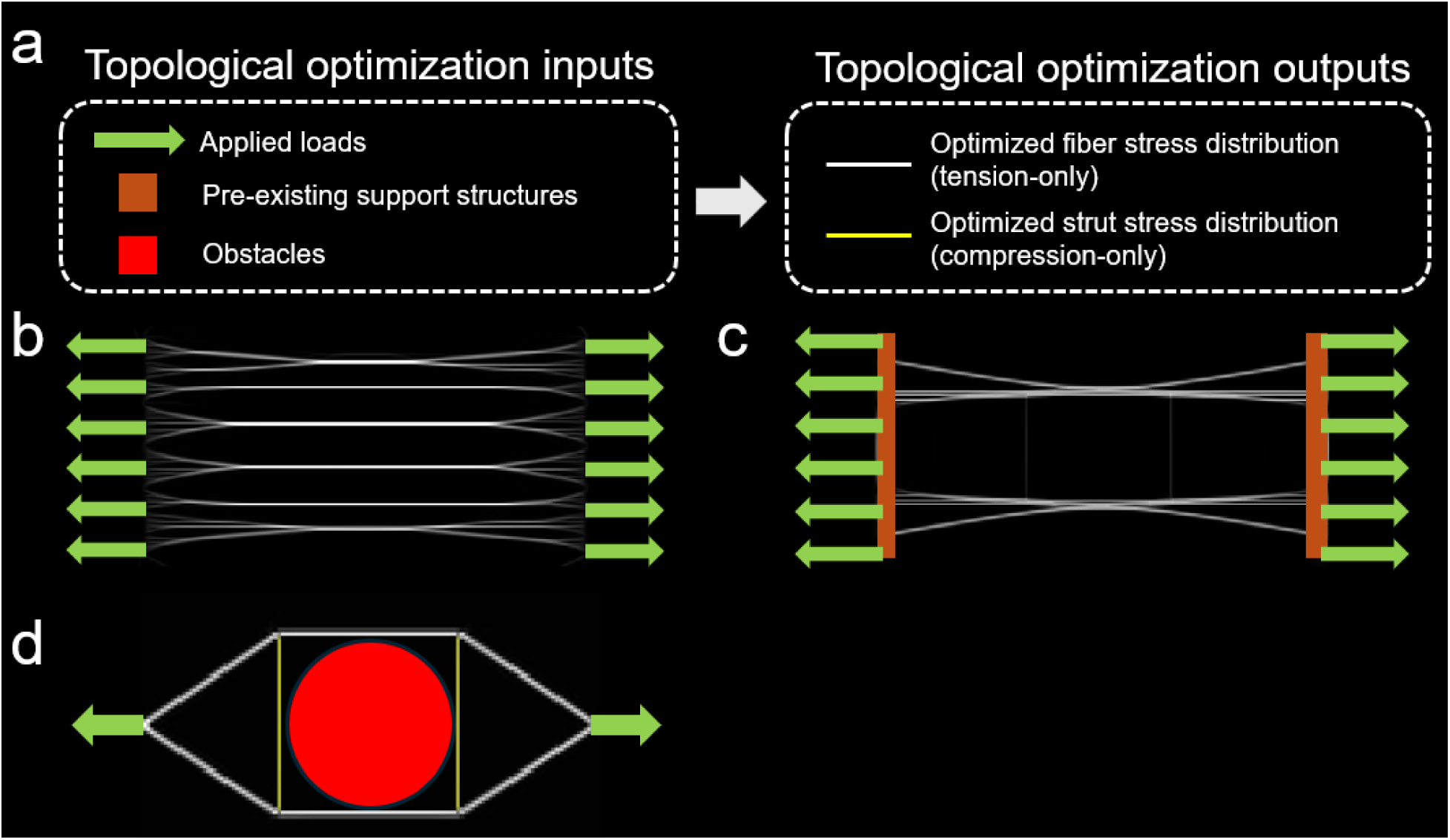
Overview of topological optimization inputs and outputs (a) with illustrative test examples (b-d). Under distributed uniaxial loads (b) the resulting fiber distribution spans the sites of force application but converges into a sparser stress set of horizontal fibers at higher stress to minimize material use. Placing pre-existing support structures at the sites of load application (c) allows for a more compact configuration with the insertion points into the structures automatically identified by the algorithm. When an obstacle is placed between loads (d) the algorithm automatically identifies new structures required to redirect loads from the obstacle. These examples, computed in our code, were inspired from examples in Rowe and Chern [24].

The output of the optimization process is a sparse stress tensor field that is consistent with the loading conditions and satisfies the other constraints. Since the cost function is designed to ensure the stress tensor has only one large eigenvalue throughout most of the domain, the trace of the stress tensor can be used to visualize the magnitude of this field. Locations where the stress is zero or negligible relative to the full scale can be interpreted as void locations. Locations where the stress is finite are interpreted as containing structural members oriented along the eigenvector(s) of the stress tensor. Rowe and Chern’s original algorithm allowed for both tension and compression members [24]. This behavior is recovered by using the eigenvalue update rule from Eq. (25) for all eigenvalues instead of only tensile members. However, unless otherwise mentioned, we only allowed tensile members.

### Topology optimization example models

To verify the implementation of the model and illustrate its behavior, we created three 2D example models. All models were solved on a 1 m x 1 m, 256 x 256 element grid with the outermost boundary elements defined as obstacles. The input for the first model was a set of two distributed 1N horizontal loads pulling in opposite directions as illustrated in Figure 1b. *σ_max_* was set to 30 N/m, *σ_cr_* was set to 1 N/m, and 500 iterations of the algorithm were performed. The input for the second model was similar except it included pre-existing structures at the sites of load application as illustrated in Figure 1c. The other algorithm parameters remained unchanged. The input for the third model was two 1N horizontal point loads in opposite directions with a circular obstacle of radius 0.1 m between them as illustrated in Figure 1d. The goal of this example was to replicate results from Rowe and Chern’s publication [24] so we allowed compressive members (struts) in this model in addition to tensile members (fibers). *σ_max_* was set to 100 N/m, *σ_cr_* was set to 5 N/m, and 500 iterations of the algorithm were performed.

### Model of the optic nerve head

We developed a topology optimization model of the optic nerve head to understand how different conditions affect optimal LC connective tissue architecture. The features described in this section were shared by all models unless otherwise noted. The optimization domain was divided into four cylindrical layers. The center of the domain was a circular vessel obstacle where no material could be placed. Surrounding that was the optic nerve canal where the cost function was computed normally to find the optimal LC architecture. This region varied in size and cross-sectional shape depending on the experiment. Surrounding the canal was a 200 μm thick sclera region obtained by morphological dilation of the canal region by a spherical structure element. The sclera was assumed to be a dense, pre-existing support structure the LC builds from. The remainder of the domain outside the sclera region was defined as an obstacle.

For all models, a distributed intraocular pressure load of 15 mmHg was applied to the anterior surface of the sclera and LC cylinders, and a retrolaminar pressure load of 7 mmHg was distributed across the posterior surface [26,27]. Additional tensile tractions were distributed along the outer cylindrical walls of the sclera. The magnitude of this tension was derived from Laplace’s law for hoop tension in the eye globe:

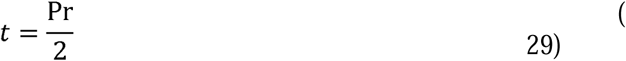

Where P is the intraocular pressure and r is the globe radius (set to 12.5 mm unless otherwise noted). The anterior-posterior component of the traction vector was set to balance the applied loads on the anterior and posterior surfaces of the model, and the component normal to the cylindrical wall was chosen to match the total magnitude from Eq. (29). A schematic representation of the model is displayed in Figure 2.

**Figure 2:**
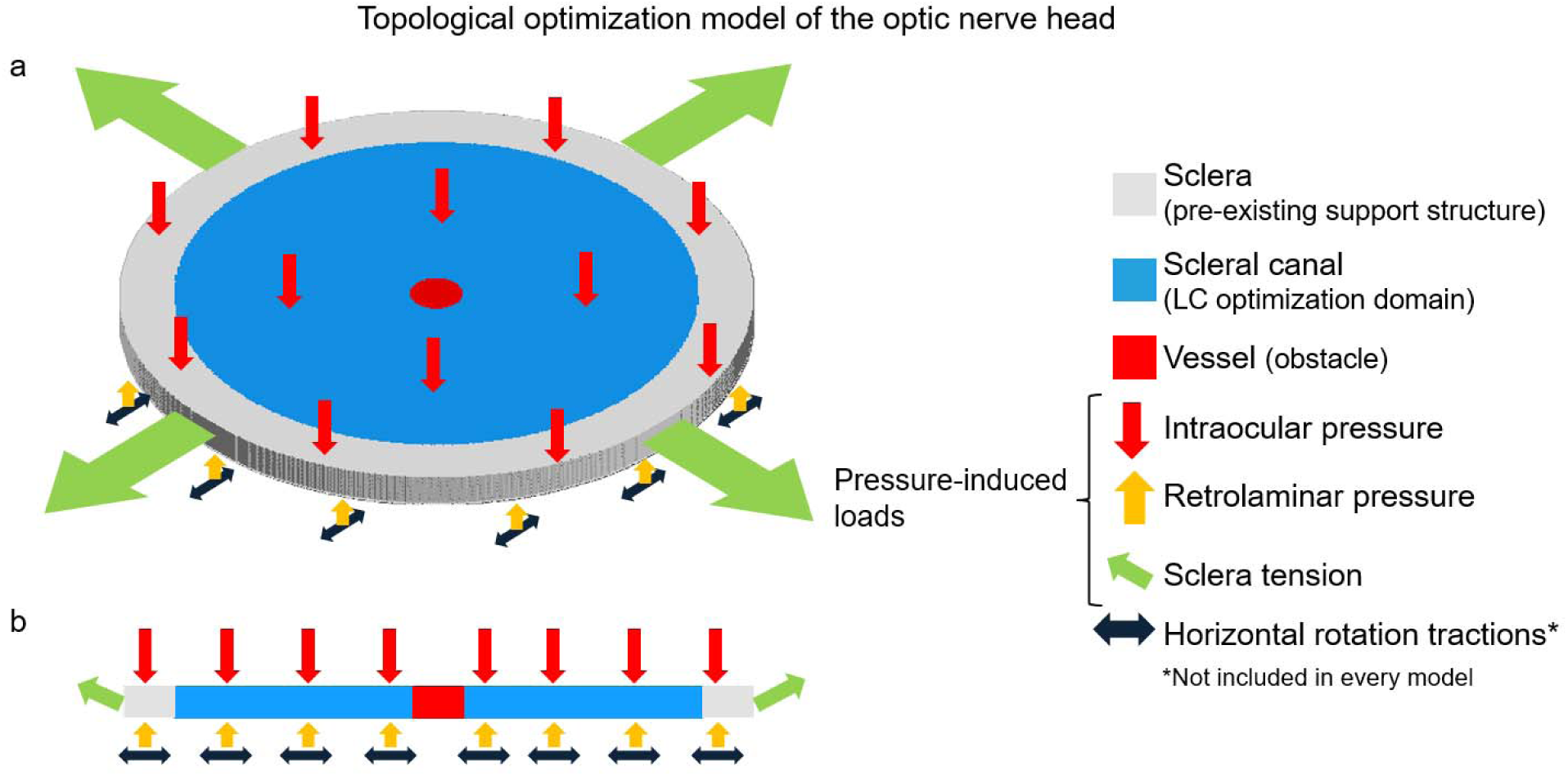
Schematic of the topological optimization model of the optic nerve head from an isometric view (a) and cross-section (b). The pressure-induced loads were applied to all models in the study and consisted of the intraocular pressure on the anterior surface, retrolaminar pressure on the posterior surface, and sclera tension on the sclera hoop surface. Sclera tension included both the reaction forces necessary to balance loads on the anterior-posterior surfaces and an outward normal traction obtained from Laplace’s law for the eye globe. Horizontal shear tractions on the posterior surface were included to simulate rotation tractions in some models, but not all. Models solved for the optimal LC fiber architecture in the scleral canal region of the domain.

Model results are visualized from the trace of the stress tensor in the optic nerve canal region (omitting the sclera region) where regions of high stress correspond to regions of high fiber material density. Volumetric data was rendered using Avizo 3D 2022.2 (Thermo Fisher Scientific, Waltham, Massachusetts, USA).

### Optimization parameter study on the optic nerve head model

To identify the role of optimization algorithm parameters we solved models with a circular optic nerve canal with diameter 2 mm. A 2.5 mm x 2.5 mm x 0.25 mm rectangular domain was discretized into 400 x 400 x 40 element voxel grid. The parameters varied were *σ_max_* and *σ_cr_*. *σ_max_* is the maximum allowed tensile stress in an element. *σ_cr_* controls the nonlinear step size along the tensile material cost gradient. A set of models was run with *σ_cr_* fixed at 70 kPa and *σ_max_* varied from 1 – 5 MPa in 1 MPa increments. Another set was run with *σ_max_* fixed at 5 MPa and *σ_cr_* varied from 10 – 90 kPa in 20 kPa increments. All models were run for 1000 iterations.

### Elliptical canal models of the optic nerve head

We modeled the optic nerve canal cross-section as an ellipse with different models varying the major:minor axis ratio from 1.0:1 (circular) to 2.0:1 with increments of 0.1 between [28–30]. For a given aspect ratio the absolute values of the major and minor axis diameters were set to preserve the area of a 2 mm diameter circular cross section. Optimization was performed on a 3.75 mm x 2.5 mm x 0.25 mm rectangular domain discretized into a 600 x 400 x 40 voxel grid. *σ_max_* and *σ_cr_* were set to 5 MPa and 70 kPa respectively. All models were run for 1000 iterations.

### Eye rotation models of the optic nerve head

In addition to protecting the nerve tissue from translaminar pressure, the LC protects the nerve tissue from tractions at the optic nerve head resulting from eye rotations [9,10]. To represent these tractions for horizontal eye rotations, we applied a distributed shear force on the posterior surface of the sclera and optic nerve canal pointing in the horizontal direction. Net traction integrated over the posterior surface varied from -15 mN to 15 mN with positive and negative values indicating a leftward and rightward shear respectively. These net tractions were comparable in magnitude to the net antero-posterior tractions from the pressure loads (about 9 mN). Horizontal tractions opposite the posterior surface shears were distributed uniformly along the outer cylindrical wall of the sclera to balance the rotational shear load.

We used a circular cross-section with a diameter of 2 mm for the optic nerve canal and a 400 x 400 x 40 voxel grid. Models were run for 1000 iterations with *σ_max_* and *σ_cr_* set to 5 MPa and 70 kPa respectively. When visualizing the structure for a given magnitude of shear traction, we superimposed the stress fields for leftward and rightward results at the same magnitude by taking the max element stress between the leftward and rightward model.

### Two-material models of the optic nerve head

Some animals, such as rats, do not have a collagenous LC, but instead have a network of fibrous actin known as the glial LC [11,16,17]. Actin fibers are more compliant than collagen fibers, suggesting that a glial LC could not protect nervous tissues from the same levels of stress as in a collagenous LC [31]. However, it is possible that a glial LC is adequate for the lower stresses of a smaller optic nerve head. To understand this distinction, we modified the tensile cost function and stepping procedure for the solution of two tensile phases: a strong material and a weak material. The weak material represented fibrous actin as seen in glial LCs. The strong material represented collagen. Instead of a single max tensile stress *σ_max_*, the algorithm enforced a weak material strength *σ_w_* and a strong material strength *σ_s_*. Strength in this case does not necessarily refer to the yield or ultimate strength of the material, but rather the amount of stress the material can sustain before its stretch begins negatively affecting surrounding nerve tissue [32,33]. Elements in the model with tensile stress below *σ_w_* are interpreted as containing weak material whereas elements in the model with tensile stress above *σ_w_* are interpreted as containing strong material. *σ_s_* is set as the absolute limit of tensile stress in the model (similar to *σ_max_* in the single material models). The cost function and updating procedure were modified to prioritize keeping stress below *σ_w_* if possible and only use strong material with stress above *σ_w_* when the problem demanded it.

Replacing the tensile cost function previously defined in Eq. (10) is the new cost function:

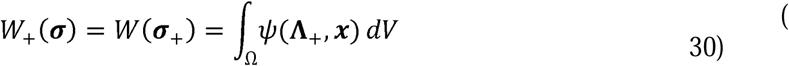

Where *ψ*(**Λ**_+_, ***x***) is the element-wise cost function in terms of ***σ***_+_ in its eigenbasis:

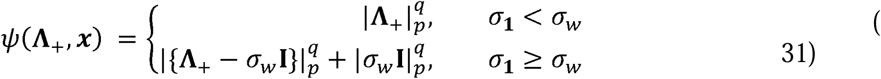

Where { } are again Macaulay brackets and *σ*_1_ is the largest eigenvalue in the element. 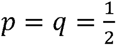 was again chosen for the cost function. The elementwise cost function for an element under uniaxial tension is displayed in Figure 3a. The cost function has a kink at *σ_w_* resulting in large gradients for stresses slightly above *σ_w_* that prevent the stress from exceeding *σ_w_* when allowed by the loading conditions. As illustrated in Figure 3b, if the loads are large enough that the stress is forced past the kink at *σ_w_* it optimizes in the strong material range preferring sparse fiber elements with stress near the new maximum allowed value *σ_s_* over many fiber elements at lower stresses.

**Figure 3:**
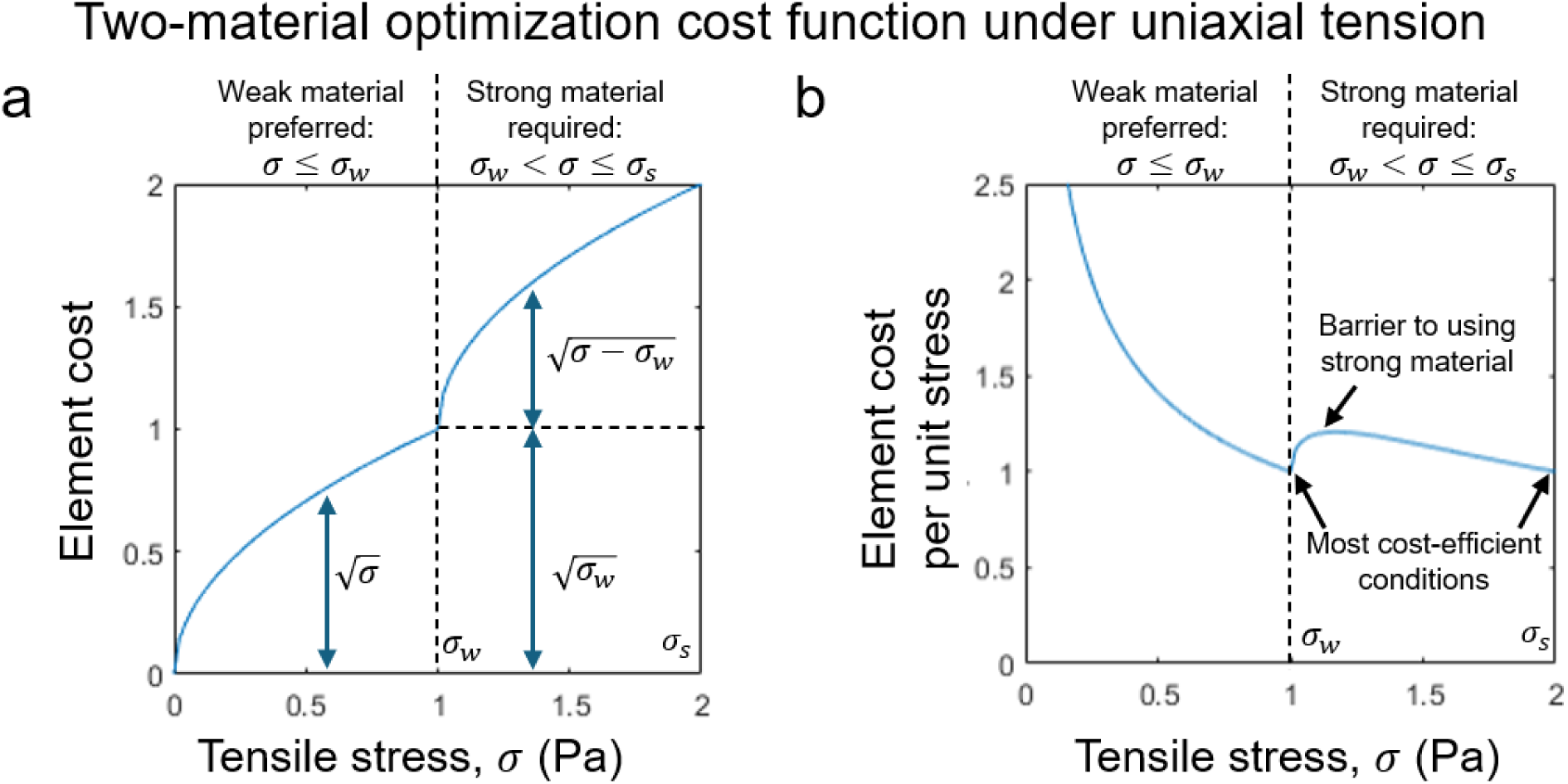
Elementwise cost for an element under uniaxial tension under the two-material cost function Eq. (31) (a). The weak material stress limit *σ_w_* and strong material limit *σ****_s_*** are shown at 1 Pa and 2 Pa respectively for illustrative clarity. The kink in the cost-stress graph at *σ_w_* results in a steep gradient that keeps the stress in the final structure at or below the weak material limit if possible. However, if the applied loads are too large the stress exceeds *σ_w_* and the algorithm will be required to use strong material. The elementwise cost normalized by stress (b) illustrates the algorithm’s preference for concentrating stress in a sparse set of elements near the weak and/or strong material limits.

Replacing the tensile eigenvalue update from Eq. (25) is the new truncated gradient descent step of Eq. (31):

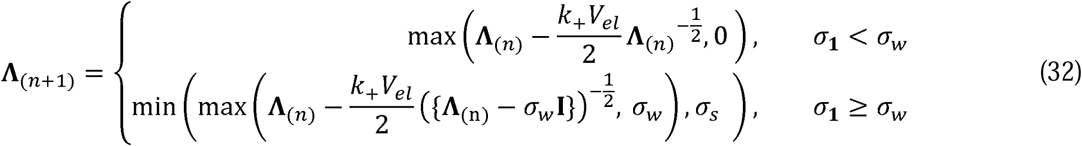

*k*_+_ is defined in terms of *σ_cr_* with Eq. (23) the same as before. The update rule behaves the same as the previous rule for weak material elements. However, strong material elements truncate their gradient descent step at *σ_w_* instead of 0 and the size of the gradient is very large slightly above *σ_w_* (more precisely, when the values of {**Λ**_(n)_ − *σ_w_***I**} are similar in magnitude to *σ_cr_*) to prevent the use of strong material unless it is necessary to satisfy the equilibrium constraint. When **Λ**_(n)_ is significantly larger than *σ_w_* + *σ_cr_* the step behaves similarly to single material optimization with strong material.

To match the geometry of a rat’s scleral canal compartment, the LC model geometry was set to the shape of a kidney bean with no central vessel obstacle (rat central vessels are located in a sperate compartment of the scleral canal separated from the glial lamina by a sheet of collagen) [34,35]. Small animal models had a globe diameter of 9 mm and a cross-sectional area equivalent to a 0.68 mm diameter circle [35]. Large animal models had a globe diameter of 25 mm and a cross-sectional area equivalent to a 2 mm diameter circle [27]. Collagen strength was fixed at *σ_s_* = 5 MPa, but actin strength was varied from *σ_w_* 100-500 kPa in 100 kPa increments. All models were run on a 600 x 600 x 40 voxel grid with *σ_cr_* = 7 kPa. Models were run for 2000 iterations since the weak material required a smaller *σ_cr_* and thus smaller optimization steps. Volumetric data was rendered using Avizo 3D 2022.2 (Thermo Fisher Scientific, Waltham, Massachusetts, USA).

### Imaging data

To compare actual biological LCs to our optimized model LCs, we obtained microscopy images from the LCs of a tree shrew, human, monkey, pig, cow and rat optic nerve heads. The images were acquired using instant polarized light microscopy (IPOL) because the technique produces clear maps of birefringent materials, like collagen and actin. In IPOL images, color encodes local fiber orientation and increased brightness can be taken as indicative of increased fiber density. Details of the IPOL technique and how brightness relates to fiber density can be found elsewhere [36,37]. Collagen has substantially stronger birefringence than actin and thus appears brighter in IPOL images. Hence, for tree shrew, human, monkey, pig, and cow we compared the models with an image of a single coronal cryosection through the LC of a given species. For the rat, however, we imaged all the sections through the optic nerve head. The images were then stacked and aligned using techniques described elsewhere [15], and a maximum intensity projection computed to compare with the models.

## Results

### Topology optimization example models

Results of three illustrative test models for the topology optimization algorithm are shown in Figure 1b-d. For the distributed uniaxial load problem (Figure 1b), fibers span the sites of load application, but merge into a sparse set of parallel horizontal fibers running between the two loads. When a pre-existing support structure is included at the sites of load application (Figure 1c), the algorithm identifies optimal points for inserting fibers into the structure and balances the load using a more compact arrangement of fibers that does not need to span the entire site of load application. When an obstacle was placed between two point loads and compressive struts were allowed along with tensile fibers (Figure 1d), the algorithm redirected fibers resulting in two compressive struts on either side of the obstacle. These results matched Rowe and Chern’s for the same model inputs [24].

### Topology optimization model of the optic nerve head

The first models of the optic nerve head had a circular scleral canal cross-section and contained only translaminar pressure loads. We examined the effects of two optimization parameters on the final architecture: *σ_max_* the maximum allowed stress in any fiber element and *σ_cr_* which controls the step size taken during the optimization procedure.

Coronal projections of the optimized architectures are shown in Figure 4. In all models a ring of fibers formed around the vessel obstacle and the remaining fibers were distributed radially from the scleral wall to that central ring. Fibers were primarily oriented radially except for a thick ring around the vessel obstacle. Holding *σ_cr_* constant at 70 kPa while varying *σ_max_* demonstrated the role of collagen strength on optimal LC architecture (Figure 4a). The central vessel ring became thinner and fewer radial fibers were required as the load-bearing capacity of the fibers increased. The differences in radial fiber density were most obvious in the range of *σ_max_* between 1 MPa and 3 MPa, but the radial fiber architecture was largely preserved beyond that range. Holding *σ_max_* constant at 5 MPa while varying *σ_cr_* demonstrated the effect of the optimization step size on results. Models with low values of *σ_cr_* were more densely covered in fibers and had a less binary distribution of tensile stress throughout the model compared to models with a large value of *σ_cr_*. Again, differences between models seemed less pronounced at the upper end of the *σ_cr_* range.

**Figure 4:**
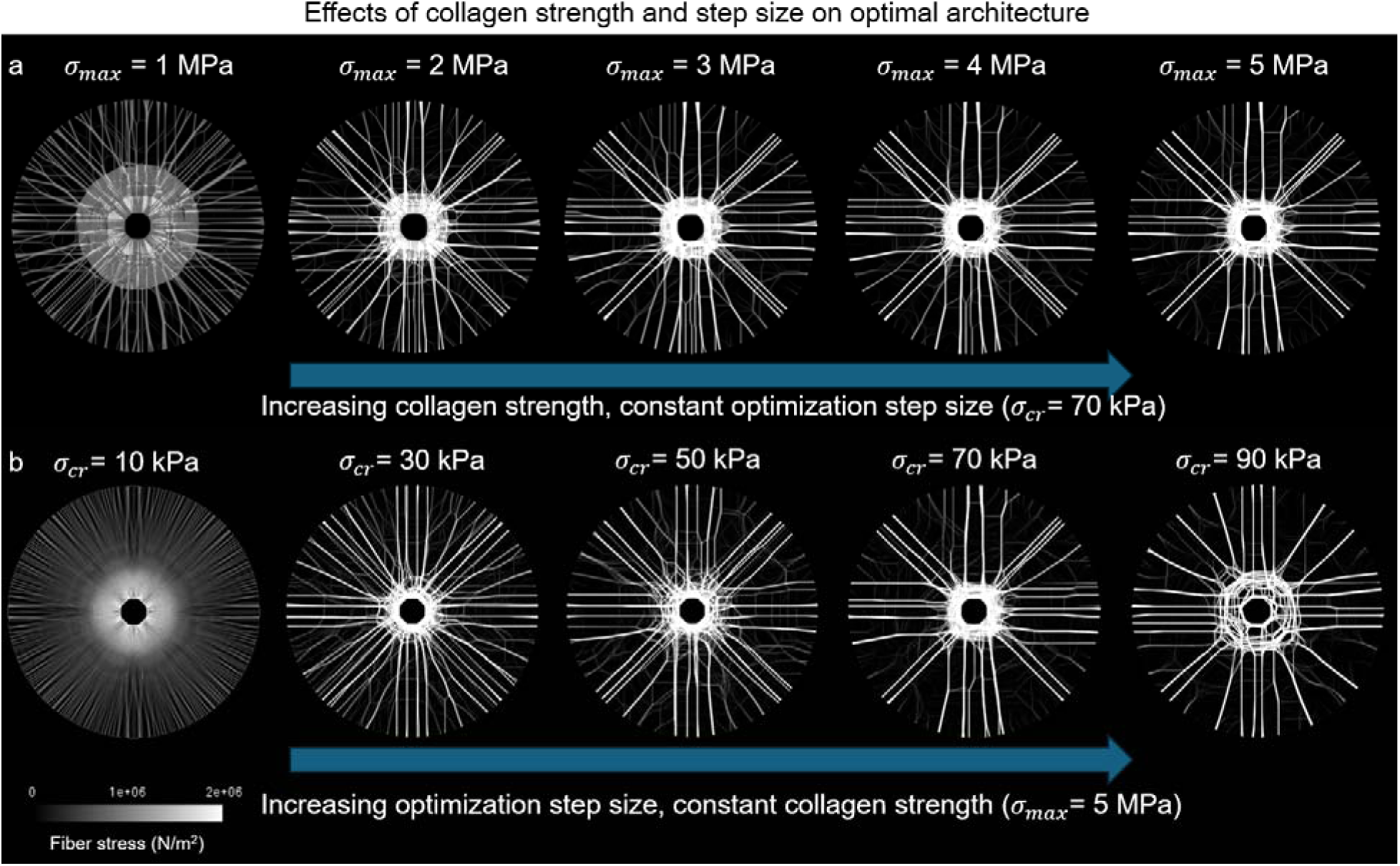
Coronal views of the fiber stress distributions optimized for translaminar pressure loads under varying model parameters using a single fiber material. Increasing collagen strength, *σ_max_*, while holding the critical stress (which is proportional to the optimization step size), *σ_cr_*, constant (a) results in sparser fiber architectures. Increasing *σ_cr_* while holding *σ_max_* constant (b) resulted in sparser architectures and less diffuse features.

**Figure 5:**
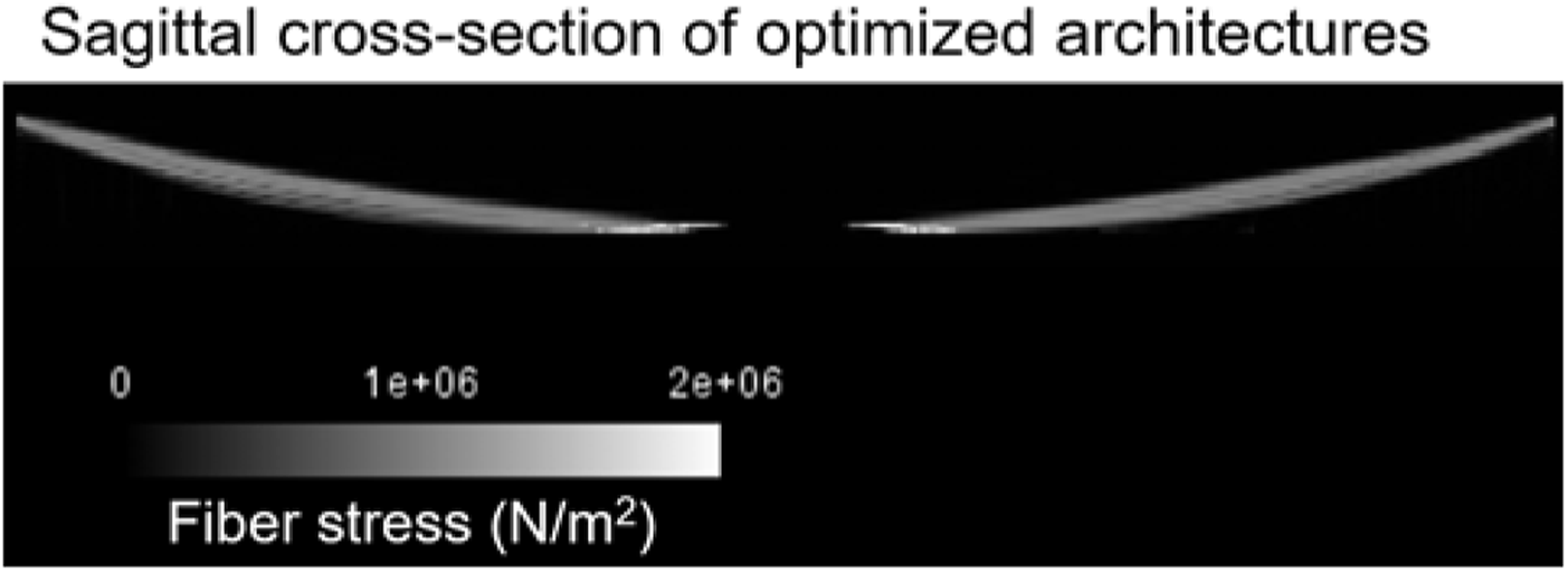
View of a 20 element (125 μm) thick sagittal cross-section taken from the center of a representative optimized fiber stress distribution (c.f. top right model of Figure 4). Fibers in all optimized architectures were similarly concentrated along a single convex surface bowing towards the posterior of the model.

From a sagittal view, all models looked like the representative case in Figure 4 of a model with *σ_max_* = 5 MPa and *σ_cr_* = 70 kPa. The LC fibers were concentrated into a single layer of elements that bowed downward in response to the translaminar pressure. The concentration of the fiber architecture into a single convex curved layer was seen in every subsequently discussed model as well and will not be explicitly mentioned for brevity.

### Effect of scleral canal geometry on LC architecture

We modeled the scleral canal cross-section as an ellipse with the aspect ratio varying from 1 (circular) to 2 (elliptical) to determine the impact of scleral canal cross-section on the LC architecture. To allow fair comparison, the cross-sectional area was preserved in each model. Coronal projections of output architectures for the elliptical canal models are displayed in Figure 6. As the aspect ratio of the ellipse increased, more of the collagen fiber architecture was oriented along the minor axis of the canal. Fibers also became sparser at the apices of the major axis where the canal cross-section’s radius of curvature was smallest. For intermediate aspect ratios, a bipolar fiber architecture formed around the vessel obstacle. An example of this bipolar architecture is indicated by the yellow arrows in Figure 6 for the model with an aspect ratio of 1.4:1. For large aspect ratios, parallel fibers spanning the minor axis of the canal dominated the architecture and the bipolar architecture became less recognizable.

**Figure 6:**
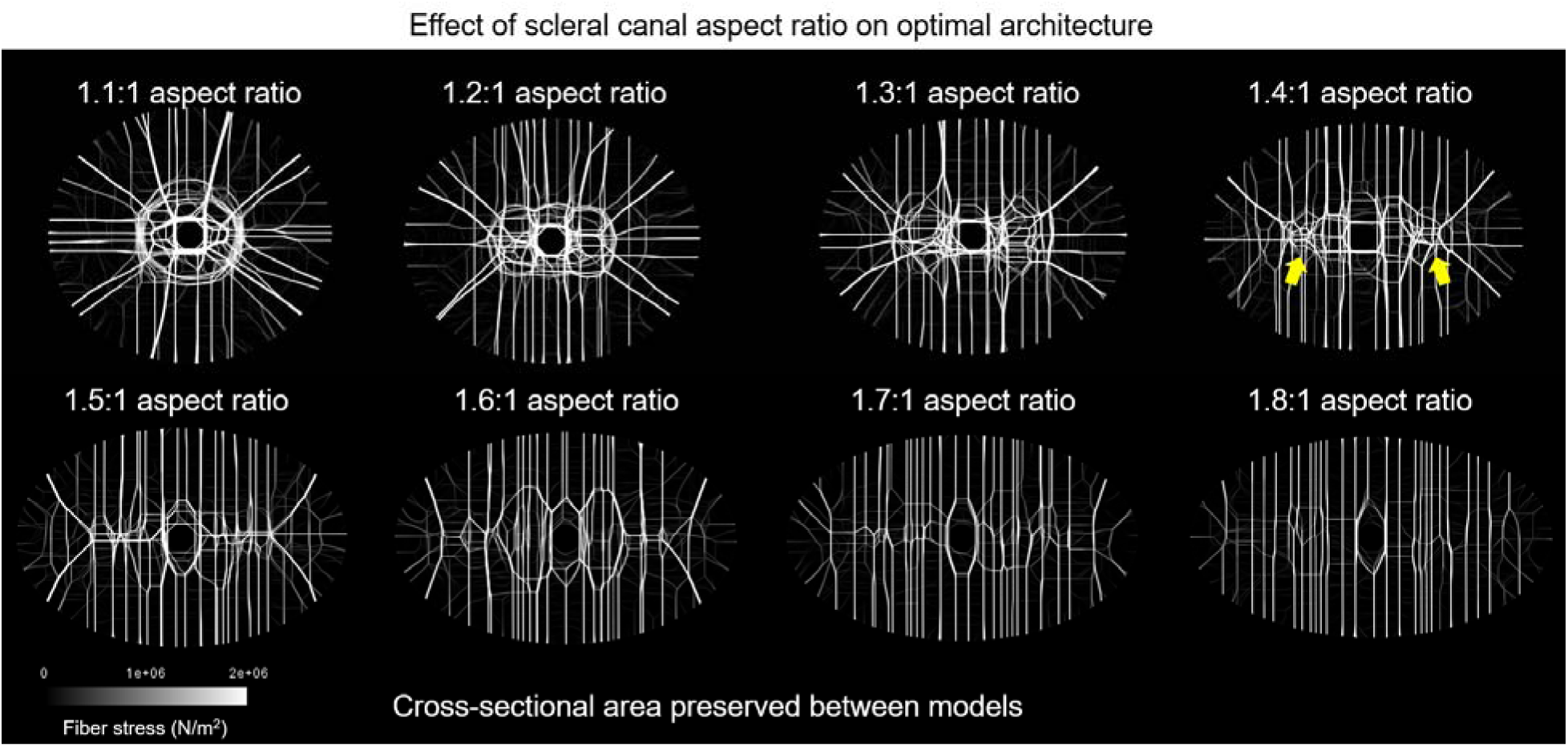
Coronal views of the fiber stress distributions optimized for translaminar pressure loads using a single fiber material when the scleral canal had an elliptical cross section of varying aspect ratio. As the scleral canal aspect ratio increases, fibers bias their orientation along the minor axis and fiber density at the periphery of the major axis decreases. For intermediate aspect ratios, fibers converge on two focal points. An example of these points is indicated by the yellow arrows. Cross-sectional area was preserved between models as the aspect ratio varied.

### Effect of eye rotation tractions on LC architecture

We applied additional shear tractions on the posterior surface of the scleral canal in addition to the translaminar pressure loads to simulate tractions from eye rotations in circular LCs. We assumed that species performing horizontal eye rotations would do this in both directions. To combine the results of horizontal tractions pointing left and horizontal tractions pointing right we took a voxel-wise max of the fiber stress between the two volumes. The volumes were then rendered in coronal view to produce Figure 7.

**Figure 7:**
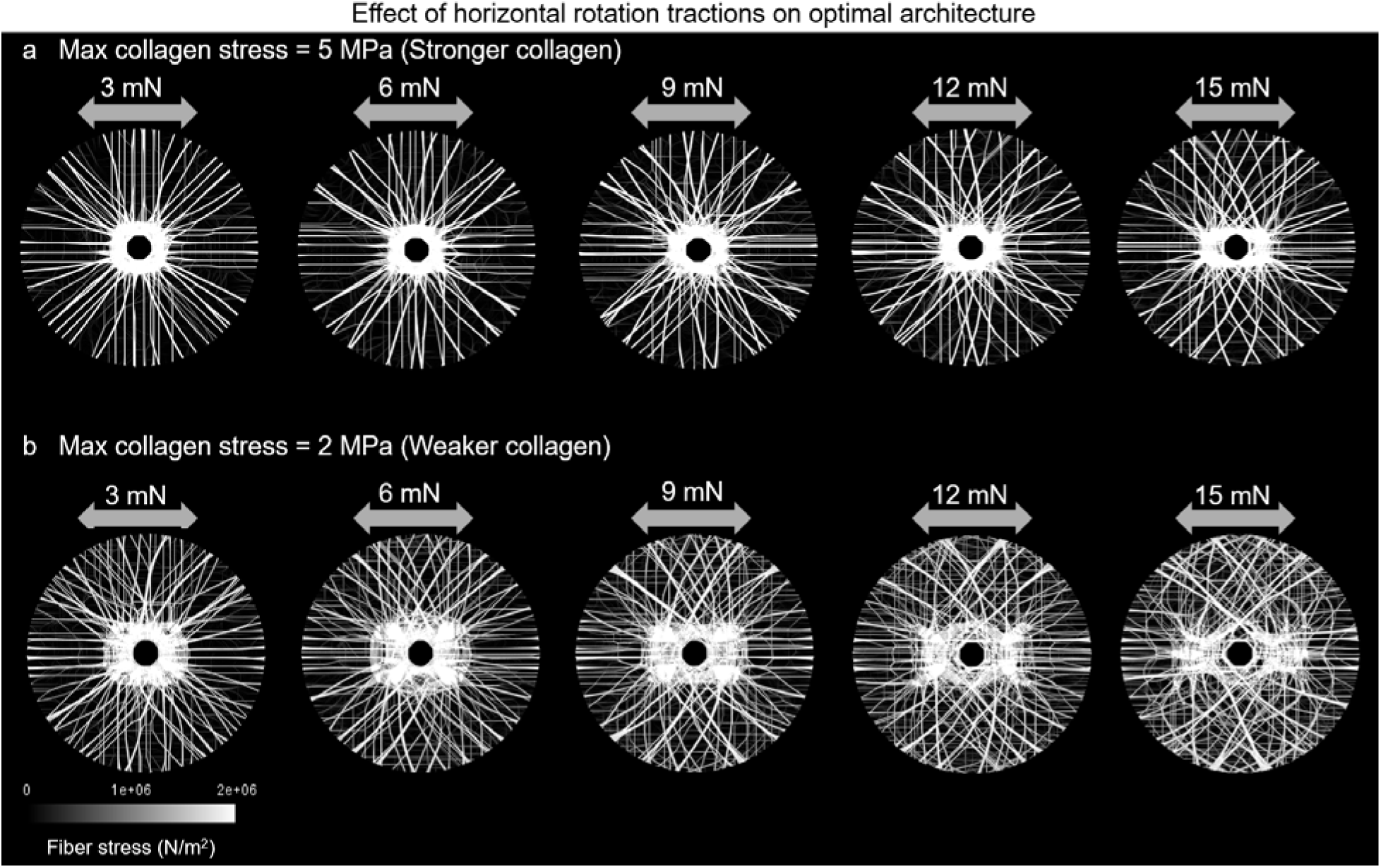
Coronal views of the fiber stress distributions optimized for horizontal rotation tractions in addition to translaminar pressure loads using a single fiber material. Results with rotations tractions pointing left were superimposed on results with rotation tractions pointing right at the same magnitude for a complete picture. For models with a higher collagen strength (a), lower traction magnitudes did not deviate much from the radial architecture, but as tractions increased fibers skewed more to the left and right to form meshwork crossings in the inferior and superior quadrants. Models with lower collagen strength (b) had a skewed architecture at smaller loads and had more complex changes in the dense collagen ring surrounding the central vessel obstacle as the traction magnitude increased.

Results for models with *σ_max_* = 5 MPa are shown in Figure 7a. Fiber architecture when horizontal rotation tractions were included varied between quadrants of the LC. In the nasal-temporal quadrants, fibers ran parallel to each other in the horizontal direction. In the inferior-superior quadrants, fibers curved left and right to bear the rotation tractions. Consequently, as the magnitude of tractions increased, the curving of fibers in these quadrants resulted in crossing meshwork patterns. The ring of collagen around the central vessel modestly elongated along the horizontal axis as tractions increased.

Additional results for models with a lower strength, *σ_max_* = 2 MPa, are shown in Figure 7b. These models displayed similar interquadrant differences to the models with stronger fibers, but with greater collagen density. Notably, the crossing meshwork patterns appeared at a lower rotation traction magnitude. When the magnitude of the tractions was 15 mN, looping structures enclosing the center of the model formed that were elongated along the vertical axis. The ring of collagen around the central vessel experienced greater elongation with the weaker models. Additionally, as the traction magnitude increased, the periphery of the ring along the horizontal axis flanged outward into more complex shapes.

### Comparison of model results to histological data

Figure 8 compares architectures derived from the topology optimization model to IPOL images of optic nerve head samples from five species. The tree shrew LC (Figure 8a) had a wheel-and-spoke architecture consisting of radial fibers directed toward a ring of collagen around the central vessels. These features were replicated in the topology optimization models with circular scleral canal cross-sections and only translaminar pressure loads applied (Figure 8b) pictured for comparison is the model with *σ_cr_* = 70 kPa and *σ_max_* **=** 2 MPa). The cow optic nerve head (Figure 8c) featured a scleral canal that was more elongated on the horizontal axis. The LC fibers biased in the direction of the shorter vertical axis and were sparser near the horizontal apices. Additionally, the LC architecture featured two dipole locations (indicated by the arrows in Figure 8c) where collagen fibers converged into regions of high density. These features from the cow LC were best replicated by the model with an elliptical scleral canal cross-section with an aspect ratio of 1.4:1 optimized for translaminar pressure (Figure 8d). The human LC collagen architecture (Figure 8e) had a bias towards horizontally oriented fibers but still had significant variation in the vertical component of fiber orientation. This feature was replicated by the topology optimization models that included rotational shear tractions along the horizontal axis in addition to translaminar pressure tractions (Figure 8h). Pictured for comparison is the model with *σ_max_* **=** 5 MPa and 15 mN shear traction magnitudes. Both the monkey (Figure 8f) and pig (Figure 8g) LC architectures had fibers distributed in crossing meshwork patterns. These meshwork crossings were also present in the inferior and superior quadrant of the models that included rotational shear tractions along the horizontal axis.

**Figure 8:**
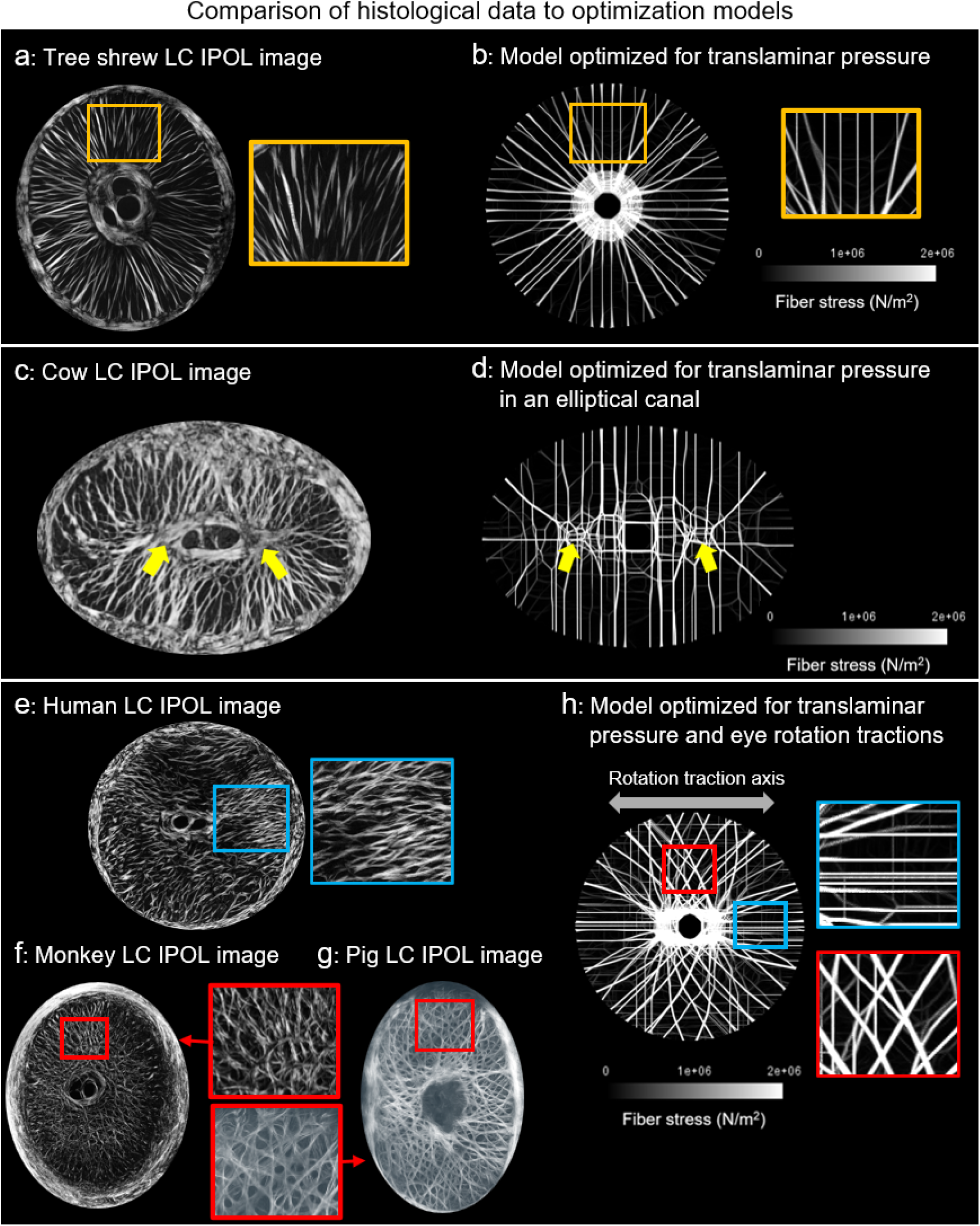
Comparisons of coronal cross sections of animal LC collagen architectures imaged with instant polarized light microscopy (IPOL) to optimization model architectures. The tree shrew LC (a) radial fiber architecture (a, inset) was replicated by the model optimized for translaminar pressure in a circular canal (b). The cow LC (c) featured a dipole architecture (yellow arrows) replicated by the model optimized for translaminar pressure in an elliptical canal. The human LC (e) had collagen fibers biases toward the horizontal direction (e, inset) and the monkey (f) and pig (g) LCs had crossing meshwork architectures (f and g, inset). The model optimized for translaminar pressure and horizontal tractions (h) replicated the horizontal bias in the nasal-temporal quadrants (h, top inset) and the meshwork crossings in the inferior-superior quadrants (h, bottom inset).

One point of contrast between the actual animal LC architectures and the topology optimization model architectures is illustrated in Figure 4c: the actual animal collagen fibers were distributed across a thin but non-trivial portion of the scleral canal’s depth, whereas the optimization models concentrated all fibers into a single layer with a thickness at the resolution limit of the discretization grid (6.25 μm) as illustrated in Figure 4c.

### Effect of animal size on LC composition

We extended the optimization algorithm to include two types of fiber material—a weak material representing actin fibers and a strong material representing collagen fibers—to determine whether animal size could explain why rats have a glial LC while the larger animal species considered in this study have collagenous LCs. To compare the load-bearing contribution of each material within a given model, the total strain energy in each material was plotted as a function of weak material strength for both small and large animal models. In both small (Figure 9a) and large (Figure 9b) animals, the strong material was the primary component (i.e. there was a collagenous lamina) when the weak material strength was at its lowest (100 kPa) and the weak material was the primary component (i.e. there was a glial lamina) when the weak material strength was at its highest (500 kPa). However, the transition point between the scenarios occurred at distinct points for the two different sizes. In small animal models, the two material components have equal contribution when the weak material strength is between 100 and 200 kPa. In the large animal models, the two material components have equal contribution at a weak material strength of 300 kPa. <u>This means that, for actin strengths between 200-300 kPa, the model predicts a glial lamina is optimal for small animals and a collagenous lamina is optimal for large animals when only the size of their eye and scleral canal is varied.</u>

**Figure 9:**
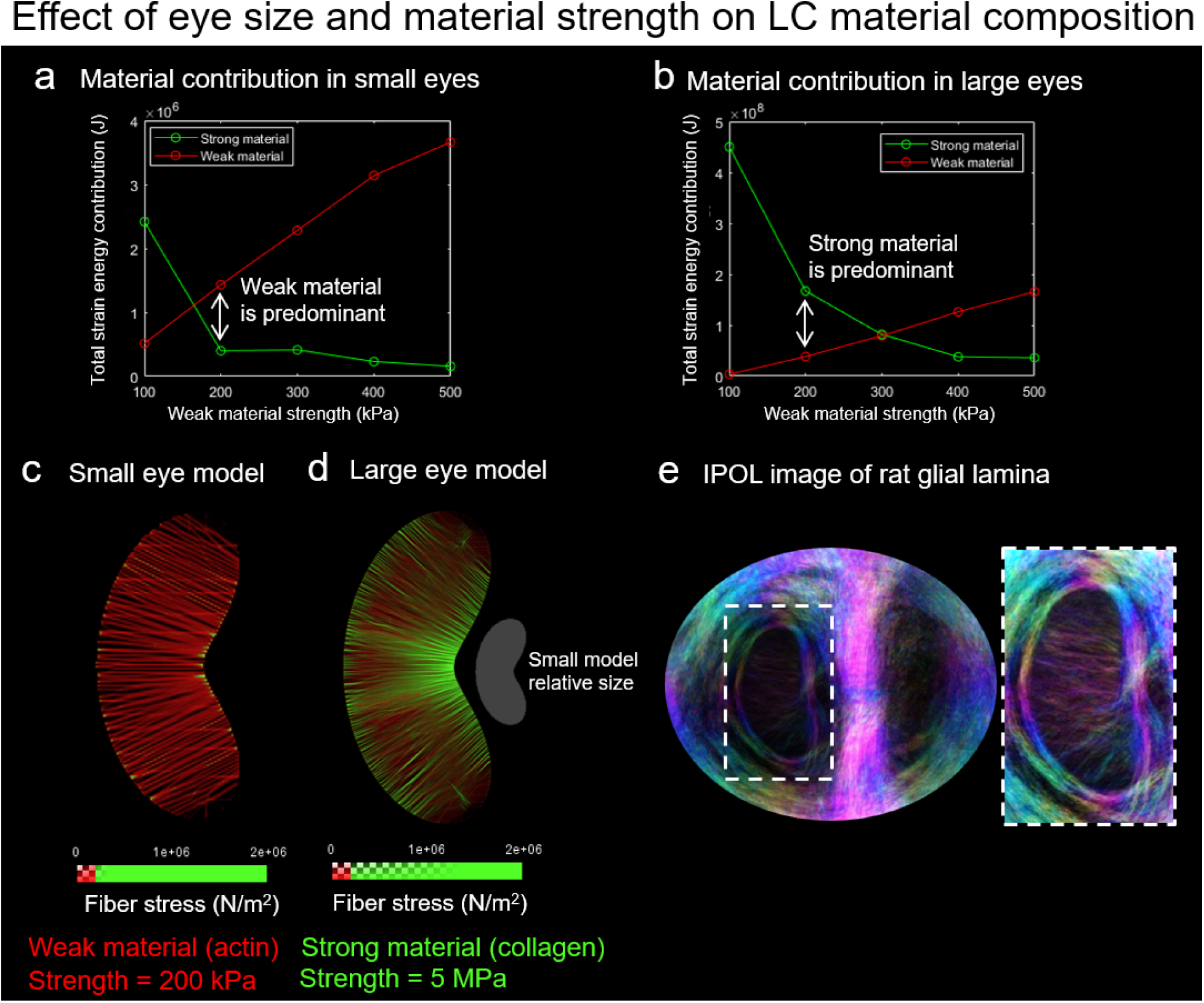
In the two material optimization models, the total strain energy contribution for the weak material increased with the weak material strength in both the small eye models (a) and large eye models (b). However, the weak material’s contribution exceeded the strong material’s contribution at a lower strength in the small eye indicating that, all else equal, a small animal could have an LC made of weaker material. Coronal views of the optimized fiber stress distributions for both the small (c) and large (d) eye models are shown for the case where the weak material strength was 200 kPa. Fibers with stress below the weak material strength (representing acting fibers composing glial lamina) were colored red and fibers with stress above the weak material strength (representing collagen fibers) were colored green. The small eye model contained only specks of collagen, whereas the large eye model had large amounts of collagen. The small eye model captured the fanning architecture of the rat’s glial lamina (e) and provides insight into the rat’s differing LC composition compared to larger animals.

Connective tissue architectures for the small (Figure 9c) and large (Figure 9d) animal models are shown when the weak material strength was set to 200 kPa. The small animal model had only specks of strong material and the weak material architecture fanned across the scleral canal in a primarily horizontal direction. The large animal model had strong material as the primary structural component though the weak material still filled some gaps in the architecture. These distinct LC compositions are consistent with literature descriptions of rat and larger animal LC compositions [11,17]. The small animal weak material architecture resembled the rat glial lamina architecture observed from IPOL (Figure 9e).

## Discussion

We set out to test the hypothesis that interspecies variety in LC architecture and composition can arise from adaptations to the unique mechanical constraints experienced by each species’ optic nerve head. To this end, we employed a novel optimization model to systematically determine how an ideal LC would adapt to the conditions we hypothesized drove interspecies differences. This allowed us to confirm whether the differences in LC architecture and composition observed between species agreed with the functions we hypothesized they adapted for. Three key results consistent with the hypothesis were obtained from this study. First, both tree shrew and cow LC architectures were replicated with only pressure-induced loads. Second, adding eye rotation tractions to models resulted in features present in human and monkey LCs. Third, our model predicted that for small animals like rats a glial lamina may be adequate to protect the optic nerve head where a collagenous lamina would otherwise be required by large animals under the same pressure conditions. Each of these findings is discussed in more detail below and the comparisons between optimization model LC architectures and actual LC architectures are summarized in Figure 8.

The tree shrew LC’s wheel-and-spoke architecture was replicated with a circular scleral canal and a translaminar pressure load. The fact that rotation tractions were not necessary to replicate the tree shrew architecture is consistent with reports that tree shrews only make small and infrequent eye rotations [38,39]. Changing the cross-sectional geometry of the scleral canal from a circle to an ellipse changed the wheel-and-spoke architecture into one reminiscent of the cow’s LC architecture. The model replicated the cow’s scleral canal elongation along its horizontal axis. Consistent with the model’s lack of rotational tractions, cows primarily track visual targets with their head rather than with eye rotations [40]. Altogether, this suggests translaminar pressure is the load of greatest importance for tree shrew and cow LCs.

Rotation tractions were required to replicate the architectures of other species. Fibers in the models with rotation tractions along the horizontal directions skewed horizontally, becoming parallel in the temporal and nasal quadrants and producing meshwork crossings in the inferior and superior quadrants. The human LC from the IPOL image had a strong horizontal bias in fibers and other studies have also observed that the primary fiber direction is along the horizontal direction albeit with significant spread in the fiber orientation [14,15]. Humans primarily navigate horizontal landscapes which could explain the biases in fiber direction seen in their LC architecture. The human eye has a more limited range of rotation in the upward direction and loses range of motion in this direction more rapidly with age than other directions [41–43]. The monkey LC had a meshwork architecture similar to the inferior and superior quadrants of the optimization model with rotational tractions. Monkeys make extensive use of fast and large eye rotations to navigate their surroundings [44]. For monkeys, vertical and horizontal rotations are both likely of importance since monkeys navigate vertical treetop environments which could explain why this meshwork architecture appears in all quadrants of their eye. The pig LC had a similar meshwork architecture to the monkey LC. Pigs share similar eye positioning and orbital anatomy to cows [45] but we could find no literature commenting on pig eye rotations in physiological conditions. Both cows and pigs have well-developed retractor bulbi muscles for retracting the eye into its socket unlike primates. Whether or not this motion induces significant tractions and deformations in the optic nerve head tissues has yet to be studied.

In addition to capturing features of how fibers are distributed in the LC architecture, the model was also able to capture differences in composition between each species using a two-material extension of the topology optimization model. Unlike the other species considered in this study which had collagenous laminas, rats have glial laminas primarily composed of actin and glial fibrillary acidic protein [11,16,17]. Glial lamina is more compliant than collagenous lamina [31] suggesting it cannot provide the same level of protection to the neural axons under high values of stress as the collagenous lamina. As an animal’s scleral canal grows in size, so does the effective bending moment the LC must resist in response to the translaminar pressure gradient. Similarly, as the globe size of an animal’s eye increases so do the hoop tensions in the sclera required to resist a given amount of intraocular pressure [27,33,46]. Thus even with the same translaminar and intraocular pressures, size plays an important role in the magnitude of the loads experienced by the LC. Our model showed that depending on the relative strength of collagen and actin components this difference is enough to drive differences in LC composition between animals.

These results came from a novel optimization methodology for understanding fiber-based architectures. Previous work by Grytz et al. also used a structural optimization approach to understand collagen fiber organization in the sclera and LC by modeling these two structures as fiber-reinforced composites and optimizing fiber orientations [47]. Our work was novel in that it optimized not just the orientation but the spatial distribution of fibers which allowed us to directly compare the pore and beam architectures of our model LCs to histological images. Our work also differed from fiber-composite approaches by seeking to capture the continuous load-bearing behavior of fibers across macroscale distances. This behavior was captured via constitutive constraints on the stress tensor that eliminated modes of stress inconsistent with the mechanics of the fibers. These constraints allowed us to better understand how an architecture capable of withstanding compressive, bending, and shear forces could emerge from structural components that function primarily in uniaxial tension. A reader may notice that our approach considered fibers but not neural, vascular, or other lamina components. By formulating the problem in this way, our intent was to find a set of fibers that effectively bore all applied loads. This implies that the fibrous architectures were sufficient to redirect loads away from other components and concentrate them into fibers specifically intended to sustain loads.

Several important parameters influenced the results of the model including the strength of the fibers, the magnitude of the applied tractions, and the critical stress which influenced both the sparsity of the final solution and the step rate of the algorithm. The fiber strength was an unknown that may vary from species to species depending on protein-level differences in fiber composition. In both the one and two-material models we showed results for a variety of fiber strengths to illustrate the impact of this unknown parameter on model results. In particular, the transition point from glial LC to collagenous LC was sensitive to the relative values of actin and collagen strength (Figure 9a and 9b). This sensitivity shows that size should not be considered the only factor that determines the composition of the LC even if it is an important factor. Another important detail to consider when interpreting the strength of the fibers is the magnitude of the applied loads. For this study, we chose to hold pressure loads constant and vary fiber strength. However, the results were ultimately sensitive to the relative values of the strength and loading magnitude. A larger critical stress resulted in sparser architectures, but the sparsity of the final architecture was also constrained by the upper bound on the stress, so the critical stress’s influence plateaued at a certain point and had less influence on the architecture (Figure 4b). A strength of this model is its ability to disentangle the influence of these overlapping factors.

### Limitations

Although the fiber architectures resulting from the topology optimization model resembled real LCs from a coronal view, there was an important difference along the depth of the fiber architecture. In actual LCs, the collagen architecture has a 3D architecture along the scleral canal’s depth [48,49]; however, the architectures produced in the model had their fibers concentrated almost entirely into a single sheet. The reason for this difference and its mechanical significance is not clear. One possibility is that unlike real collagen, the modeled fibers can be placed arbitrarily close to each other allowing the full structure to be compressed to a single layer whereas the biological processes producing collagen fibers may have space constraints (e.g. on the size and shape of bundle cross-sections) that require spreading out bundles of collagen. Our model did not include every possible biological constraint or feature that could influence LC architecture and composition which may result in some inaccuracies.

Eye rotation tractions were simplified to uniform shear tractions across the posterior of the lamina and sclera. Previous work has emphasized the tractions between the optic nerve sheath and the peripapillary sclera that arise during eye rotations [9,10]. The LC and sclera distort in response to these sclera tractions which induces further tractions between the LC and nerve tissue crossing through it. The precise magnitude and distribution of these tractions depend recursively and nonlinearly on the architecture and geometric configuration of the LC and sclera which the current model did not account for. The choice of a uniform shear distribution across the posterior sclera and LC surface is a simplification, but one that ensures the resulting architecture provides coverage to the entire LC-optic nerve interface and that captures an aspect of loading distinct from pressure loading.

The optimization formulation we used solved for a single loaded configuration of the fiber architecture. However, the magnitude and type of loading the LC experiences can vary over time [50,51]. Accounting for this variety of loads poses two main challenges: 1) Incorporating the non-linear influence of different loads on the fiber architecture’s geometric configuration (and vice versa) which is highly dependent on the material properties of the fibers. This includes non-linearities such as fiber (un)crimping [22,23] and fiber-fiber contact [52] which can impact the mechanical response of the LC under different loading conditions. 2) Determining how each of the many possible loading conditions contribute to the optimization problem’s overall cost and constraint formulations. For our horizontal rotation results we superimposed results from leftward and rightward tractions to get a more complete picture of what an architecture accounting for both loads would look like, but an optimization formulation that considers both loading constraints simultaneously may find a more efficient architecture that satisfies both loading constraints.

Collagen fiber placement is likely constrained by a variety of other biological factors not included in the model such as sharing space with important functional components in the LC like smaller blood vessels [53] and nerve axons [54], additional uses of collagen such as providing a scaffold for angiogenesis [55], and the specific metabolic and biotransport constraints of collagen assembly [56]. The relative importance of each of these constraints on the final collagen architecture compared to the constraints modeled in this study is not obvious. Without these biological processes the model presented here is not intended to be a model of how the LC collagen architecture forms or remodels. Instead, the model was used to determine the ideal architecture for mature LCs under a given set of mechanical conditions. The fact that this was adequate to capture notable features from various animal species suggests that, whether intergenerationally through evolutionary pressure or via complex reactive growth dynamics throughout an individual’s lifetime, adaptation to mechanical loads may play an important role in determining the connective tissue architecture and composition of the LC.

Additional limitations arise from the numerical optimization framework. Our choice of cost function and cost function coefficients was not unique in their ability to produce sparse fiber architectures. For this study, we chose to vary the parameters most directly tied to physiological interpretation rather than explore the wide landscape of potential cost functions; however, other choices may potentially result in more accurate results. The gradient descent-based update scheme used to solve the problem may be susceptible to local minima. Results provided in the supplemental materials Figure S1 indicate that although individual fibers and beams may have been influenced by these local minima, the broader architecture was robust to variations in initial state. We prioritized easy repeatability and stability for our initialization strategy while noting that the mechanisms driving adaptation in each species’ LC could also produce local minima.

Readers should be aware that we compare model results to only a single specimen of each species. Our experience is that the examples shown capture the essence of each species, but we must acknowledge that individual morphology may not be representative of a species as a whole.

## Conclusion

This study introduced a novel topology optimization framework for understanding the specializations of LC fiber architecture and composition in different species. By varying only the mechanical loads and scleral canal geometry of the model, distinct features from different species’ LC architectures were replicated. The mechanical distinctions between models agree with how each species uses its eyes to navigate its environment. Extending the model to include two fiber materials captured how differences in size may drive the differences in fiber composition between rat LCs and larger mammal species.

## Acknowledgements

This research was made possible by funding from the Canadian Institutes of Health Research, NIH R01-EY023966, R01-EY031708, P30-EY008098 and T32-EY017271 (Bethesda, MD), Research to Prevent Blindness (unrestricted grant to UPMC ophthalmology, and Stein innovation award to Sigal IA), and the BrightFocus Foundation.

Special thanks to Dr. Rafael Grytz at the University of Alabama Birmingham for the tree shrew eye sample.

## Algorithms

### Algorithm 1

Topological Optimization of Fiber Architectures

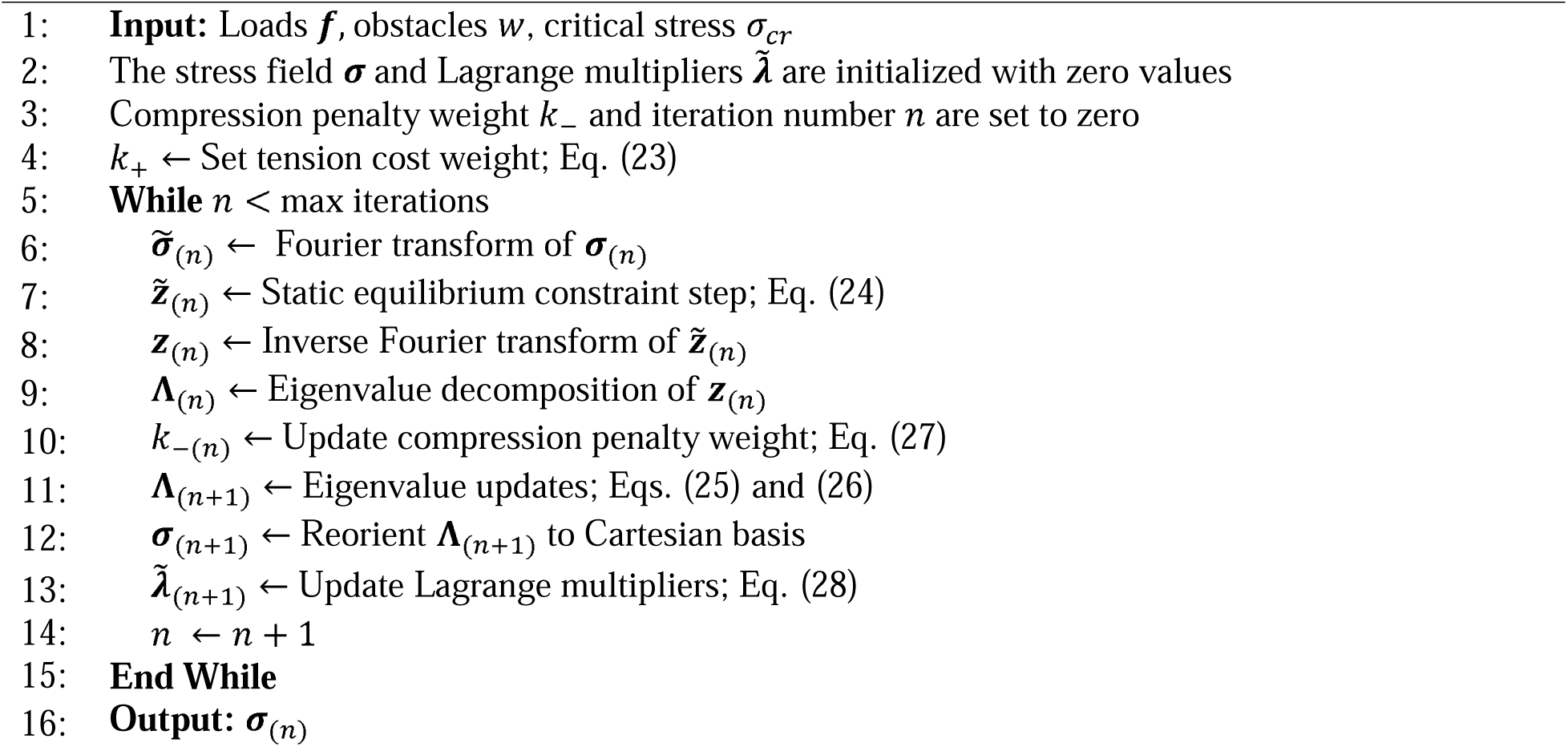

