## Supplementary material for "Interspecies differences in lamina cribrosa fiber composition and architecture might arise from differences in optic nerve head size and mechanical constraints": Figure S1

**Effect of initial state on model output**

Gradient-based optimization methods are susceptible to local minima and the choice of initial state for the optimization variables. To understand the influence of the initial state on the final solution for our models, we applied random perturbations to the initial state of an optic nerve head model with circular scleral canal and translaminar pressure loading conditions (cf. Figure 8b for unperturbed results).

In more detail, the initial state of the stress tensor in the perturbed models was generated from a Gaussian random field (GRF). Gaussian random fields are obtained by adding together many randomly oriented sine waves of fixed wavelength (in this case 1000 waves) on the voxel domain. Three models were perturbed with a wave number of 15 (i.e. the wavelength was 1/15^th^ domain size) and three models were perturbed with a wave number of 30. The values of all GRFs were re-scaled to the range -100 kPa to 100 kPa. The stress tensor was then initialized to a hydrostatic pressure scaled by the GRF at each voxel location. For all six perturbed models $\sigma_{cr}$ was set to 70 kPa and $\sigma_{max}$ to 2 MPa. All other model parameters were set according to rhe models in the *Methods: Optimization parameter study on the optic nerve head model* subsection.

Results from the six models compared to the unperturbed model are presented in Figure S1. Individual fibers and beams differed significantly between models even when models shared the same GRF wave number. However, all perturbed models retained the same wheel and spoke architecture of the unperturbed models. Note that the perturbations to the initial state were intentionally given a large amplitude and frequency to test the model’s response to variations in initial state. These perturbations do not necessarily reflect any physically meaningful process or source of variance in the sampled species.

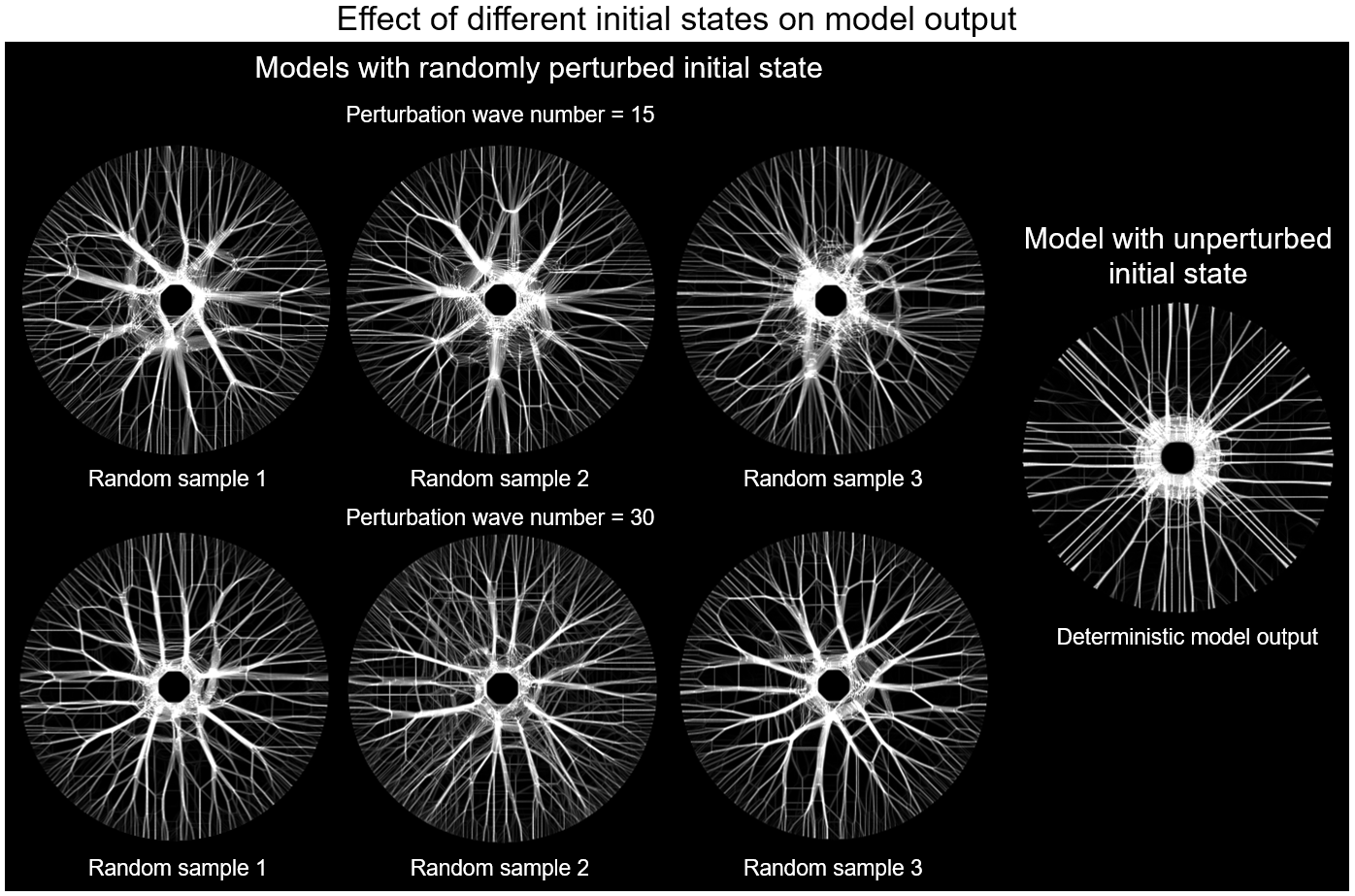

Figure S1: Effect of different initial states on model output. Six models were perturbed with a randomly sampled Gaussian random field (GRF) upon initialization (left) but were otherwise identical to the model optimized for translaminar pressure in a circular canal (right, cf. Figure 8b). Three of the perturbed models used GRFs with a wave number of 15 (top left) and three used higher frequency GRFs with a wave number of 30 (bottom left). Although individual fibers and beams differ significantly from the unperturbed model, the wheel and spoke architecture was preserved with the initial perturbations.
